# mTOR dependent transformed human cells have a distinct set of essential genes from bcr-abl transformed cells

**DOI:** 10.1101/737817

**Authors:** Don Benjamin, Marco Colombi, Christoph Moroni, Michael N. Hall

## Abstract

Constitutively active intracellular signaling drives and sustains cancer growth. The mTOR kinase integrates multiple inputs sensing nutrient, energy and growth factor levels to promote protein synthesis and anabolic metabolism, and is hyperactivated in a broad range of cancers. The bcr-abl kinase is a fusion protein generated by chromosomal translocation and gives rise to chronic myeloid leukemia and a small sub-set of leukemias. Using an in vitro transformed murine cell model, we performed shRNA knockdown of 25 genes previously shown to be essential for mTOR dependent oncogenic growth and survival. None of the genes were essential in the bcr-abl transformed line. Interrogation of this gene set in human cancer cell lines revealed that many of these genes were essential in cells dependent on mTOR signaling (defined by sensitivity to pharmacological mTOR inhibition). However, none of the genes were essential in bcr-abl transformed K562 cells that are insensitive to mTOR inhibition. Thus there is a clear divide between cells transformed via bcr-abl directed oncogenesis and other modes of transformation where mTOR, due to its central role in regulating cell growth and metabolism, is recruited as part of an oncogenic program. These validated hits represent a set of genes essential for executing critical functions downstream of mTOR and may be novel therapeutic targets for cancer.

## Introduction

The mTOR signaling pathway is an important regulator of cellular homeostasis. By integrating inputs pertaining to nutrient, growth factor and energy levels, its output determines if a cell engages in anabolic processes (protein, nucleotide or lipid synthesis) under favourable conditions, or catabolic processes such as autophagy under adverse conditions [1]. mTOR is a serine/threonine kinase that forms two distinct multi-protein complexes: the raptor containing mTOR complex 1 (mTORC1) and rictor containing mTORC2. mTORC1 is regulated by nutrient availability and broadly controls translation, autophagy and metabolic processes. mTORC2 meanwhile, is growth factor regulated and via an important substrate, the Akt oncogene, controls cell survival.

Due to its critical role at the intersection between cell growth, metabolism and proliferation, dysregulated mTOR signaling is a frequent event in cancer [2]. Mutation of mTOR regulators such as TSC1/TSC2 or PTEN (that negatively regulate mTORC1 and mTORC2 respectively) are common oncogenic events that result in constitutive mTOR activity. Constant mTOR signaling provides support for cellular hyper-proliferation from the increased synthesis of required proteins and cellular building blocks.

We previously described an mTORC2 addicted cell model where murine mast cells are transformed by constitutive mTORC2 signaling following mutagenic knock-out of PTEN [3]. As a consequence these cells are exquisitely sensitive to mTOR inhibition compared to the parent cell line. A genome-wide shRNA screen identified genes that are conditionally essential for growth and survival only under an mTORC2-driven oncogenic program. These hits are thus candidate genes for effector proteins that are responsible for executing essential functions upon mTORC2 hyperactivation.

To confirm these hits, we re-screened the top 25 hits in an isogenic cell panel. Two independent PTEN deficient lines were derived from the parental cell line. In addition, the bcr-abl oncoprotein was used to transform the parental cell line via an alternative oncogenic pathway. The screen confirmed that these genes were also essential in the newly generated PTEN-/mTORC2-addicted cell models. Strikingly none of the genes were essential in the bcr-abl transformed line.

Here, we extended the study to a panel of human cancer cell lines. Two PTEN negative cell lines with high basal mTORC2 activity (OPM2, Jurkat) were selected, a PTEN positive leukemia cell line HL60, and a bcr-abl transformed CML cell line (K562). shRNA screening of the human counterparts of the hits previously validated in the murine cell line panel revealed that a substantial fraction were also essential for survival in the human cell line panel, with the exception of K562 where shRNA knockdown of these genes had little to no impact on growth and survival. Pharmacological inhibition of mTOR also had the least effect on K562 compared to the other cell lines, indicating that mTOR is not as critically required in bcr-abl mediated transformation. These results show that bcr-abl mediated transformation is distinct from oncogenic programs where mTOR is constitutively activated (as in the case of PTEN loss), or has been recruited to support cancer cell growth.

## Results

### In vitro transformation of mast cells

We previously described an in vitro transformation system based on murine PB3c mast cells [4]. Parental 15V4 cells are dependent on exogenous interleukin-3 (IL3) for in vitro growth and survival. These cells can be transformed to acquire growth factor independence, a hallmark of cancer, by various routes [3–5]. Transformation to IL3 independence can be achieved by loss of *Pten* expression via mutagenesis (6.5 and 6.8 cells), and is phenocopied by stable shRNA knockdown of *Pten* (shP) (Fig 1A). The Pten-cell lines have hyperactive mTORC2 signaling as measured by phosphorylation of Akt at serine 473 (Fig 1B). Although mTORC1 activity is also increased in the Pten deficient cell lines (at threonine 389 of the mTORC1 target S6K), we previously showed that these cells are addicted to mTORC2 and not mTORC1 signaling for growth and survival [3]. An alternative route of in vitro transformation in this system is expression of the bcr-abl oncoprotein in 15V4 cells to generate IL3 independent b-abl cells (Fig 1A) [6]. Unlike the Pten deficient cell lines, b-abl cells have similar basal levels of mTORC2 activity as the parental cells and a slight increase in mTORC1 signaling (Fig 1B).

**Fig 1.**
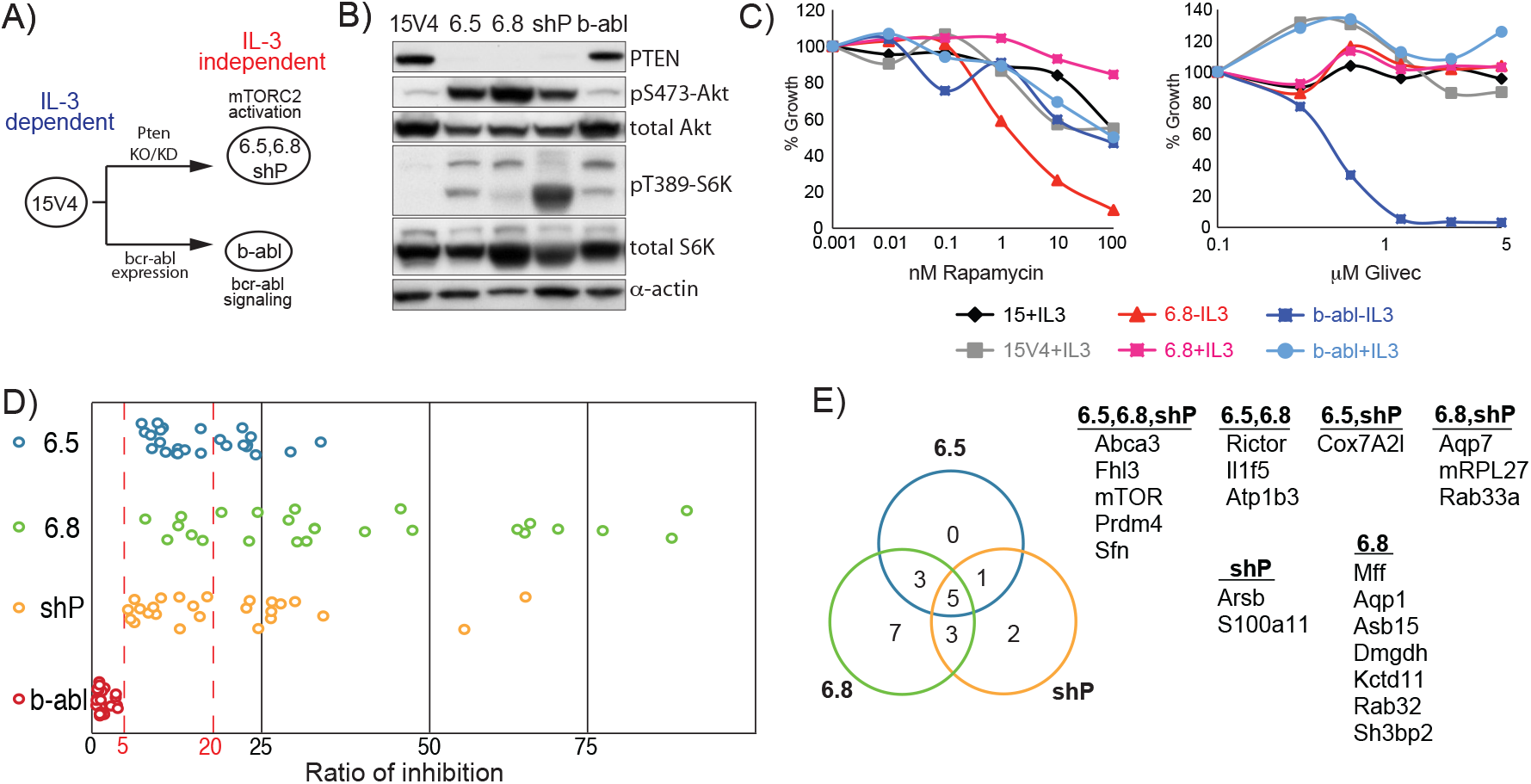
(A) Flow scheme of the in vitro transformation from IL3-dependent parental cells to IL3-independence. (B) Immunoblot showing constitutively active mTORC2 in PTEN-but not in b-abl transformed cell-lines. (C) Growth assay for cells treated for 3 days with rapamycin or Glivec. Growth was normalized to vehicle treated controls. Each data point was performed in triplicate and error bars are SD of the mean. (D) Inhibition of proliferation in the murine mast cell panel after shRNA knockdown of selected genes. The ratio of inhibition (RI) was calculated using the mean of the two best performing shRNAs and was a ratio of the growth^−IL3^/growth^+IL3^. (C) Venn diagram of genes having an RI>20%.

In these cells, IL3 signaling remains intact despite having lost their requirement for IL3. 6.8 cells are sensitive to rapamycin, as anticipated due to its addiction to mTORC2 signaling (Fig 1C). However rapamycin sensitivity is rescued by exogenous IL3, indicating that loss of mTORC2 signaling can be compensated by stimulation of IL3 signaling. Similar results were obtained with 6.5 and shP (data not shown). b-abl cells however, are sensitive to the bcr-abl inhibitor Glivec in the absence of IL3 and are rescued by exogenous IL3.

In these titrations, b-abl cells are rapamycin insensitive while the 6.8 cells are Glivec insensitive arguing that different downstream signaling pathways are activated to confer IL3 independence. Nonetheless both loss of mTOR or bcr-abl signaling can be compensated by IL3 signaling.

### Genes essential in mTORC2 addicted cells are not required in bcr-abl transformed cells

The ability to rescue cell lethality with exogenous IL3 allows identification of genes whose knockdown is conditionally lethal only in the absence of IL3. This was previously exploited to identify genes required for mTORC2 driven IL3 independent growth in 6.5, 6.8 and shP by a genome-wide shRNA screen [3]. To see if any of these genes are shared in common in bcr-abl mediated IL3 independence, we selected 25 of the highest scoring genes for screening in b-abl (the screen was repeated for 6.5, 6.8 and shP). For each gene, cells were infected with a lentivirus bearing the respective targeting shRNA (typically 5 different shRNA contructs per gene) in the presence or absence of IL3, and growth measured 5 days post-infection. Thus for each shRNA construct, a ratio of inhibition (RI) could be calculated by dividing proliferation_+IL3_/proliferation_−IL3_ (normalized to infection by a nontargeting shControl virus). The mean RI of the two highest scoring shRNAs was taken for each gene (Table S1).

The mean RI of the 25 genes are plotted for all 4 cell lines in Fig 1D. The RI’s for the mTORC2 addicted (6.5, 6.8 and shP) and b-abl cells form two distinct non-overlapping classes. Strikingly, shRNA knockdown of none of the genes has an effect on b-abl viability suggesting that they are not involved in bcr-abl driven IL3 independence. We set an arbitrary cut-off and analyzed genes having an RI greater than 20% (Fig 1E). Expectedly, as these are mTORC2 addicted cells, mTOR is a hit in all 3 cell lines while the mTORC2 specific component Rictor is also a hit in 2/3 cell lines. Several genes (mRPL27, Cox7A2l, Mff, Dmgdh) have mitochondrial functions, and as previously described, mitochondrial activity is critical for the mTORC2 addicted cells[3]. However, the remaining genes form a seemingly heterogeneous group that may nevertheless share a common role as being downstream effectors of mTORC2 signaling.

### A substantial fraction of genes essential in murine mTORC2 addicted cells are also essential in human cancer cell lines

To extend these findings from an in vitro transformed murine cell model to human cancer, we selected known PTEN negative cell lines OPM2 (multiple myeloma) and Jurkat (T-cell leukemia) as these cells have high constitutive mTORC2 activity. K562 (chronic myelogenous leukemia) expresses the bcr-abl oncoprotein and was chosen to see whether it would represent an outlier with respect to the other cell lines, similar to the b-abl cell line in the murine cell panel. HL60 (acute myeloid leukemia-M2) is PTEN positive and does not express bcr-abl, it is however insulin dependent for in vitro growth indicating a requirement for mTORC2 signaling [7]. Basal mTOR activity of these cell lines under nutrient-replete conditions (similar conditions as in shRNA screens) was determined by immunoblotting. PTEN negative OPM2 and Jurkat cells as expected had high mTORC2 activity, and slight elevation of mTORC1 signaling (Fig 2A).

**Fig 2.**
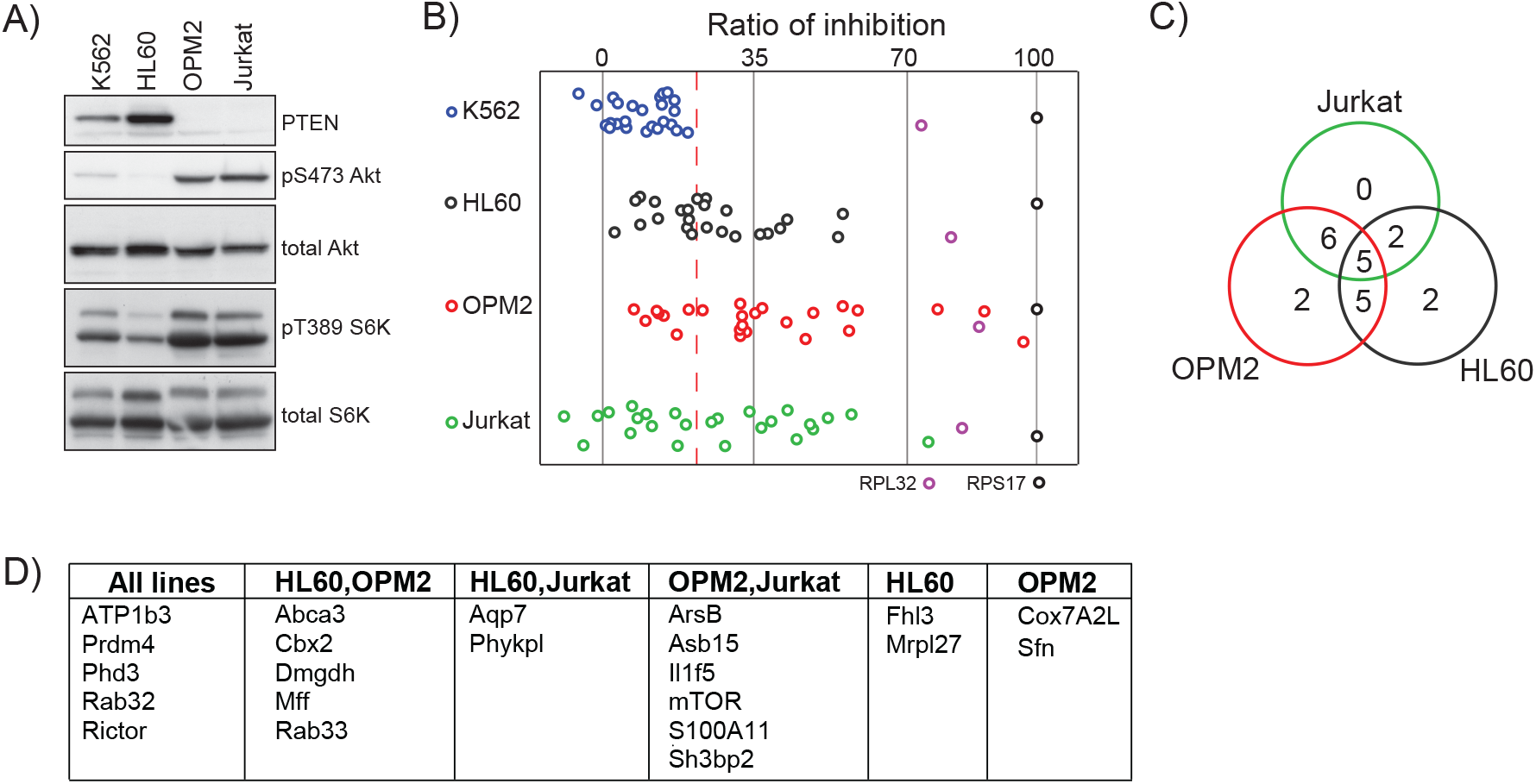
(A) Immunoblot for read-outs of mTOR activity in the human cancer cell line panel. (B) Inhibition of cell proliferation in the human cancer cell panel after shRNA knockdown of the human counterparts of the selected hit-list genes. Essential ribosomal genes RPL32 and RPS17 were positive controls for knockdown lethality. The ratio of inhibition (RI) was calculated using the mean of the two best performing shRNA constructs per gene normalized to RPS17 knockdown. (C) Venn diagram of genes having an RI>19%. (D) List of genes from (C).

Lentiviral particles for the human counterparts of the 25 essential genes in the mTORC2 addicted murine cells were obtained and used to screen the human cell panel. Unlike the previous screen, there was no possibility of growth factor stimulation corresponding to IL3 to rescue the knockdown of an essential gene. Therefore we needed to measure the infectious titer of each virus in order to discern if an effect on proliferation is biologically relevant due to the knockdown of its target gene, or a technical artefact due to low viral titer. To this end, we infected mouse bcr-abl cells with the human targeting lentiviruses and selected for puromycin resistance. The level of proliferation (as denoted by acquired puromycin resistance) was taken as a measure of the viral titer and was used to normalize infectivity of all the viruses (Table S2). shRNA’s targeting the essential ribosomal genes RPL32 and RPS17 served as positive controls for cell death. Knockdown of both ribosomal genes were lethal, with RPS17 being more effective and was used to set a baseline for cell death (Table S2).

Reduction in cell proliferation after shRNA knockdown of the 25 genes in the human cell panel is shown in Fig 2B. The ratio of inhibition for each lentivirus was calculated from (growth in infected human cells)/(growth in infected b-abl cells) and normalized to the RI of RPS17. The RI for each gene was the mean of the two best lentiviruses. shRNA knockdown of a substantial fraction of the genes inhibited proliferation in all the cell lines with the exception of bcr-abl transformed K562. This parallels the results seen in the mouse panel where only the b-abl line was unaffected by shRNA knockdown of the corresponding murine genes. Accordingly, we set a threshold for effectiveness above the K562 cluster (corresponding to a reduction in proliferation greater than 19%, red dashed line). In the non bcr-abl transformed lines, knockdown reduces cell proliferation in around half of the genes targeted (Fig 2C). Rictor knockdown was positive in all 3 cell lines and mTOR in 2 out of 3 lines (Fig 2D). Genes encoding for mitochondrial function (MRPL27, Cox7A2l, Mff, Dmgdh) were once again essential.

Thus the list of 25 essential genes obtained from the isogenic mTORC2-addicted murine screen translates very well to a heterogeneous panel of human cancer cell lines. Some of these hits, in particular those from the PTEN negative OPM2 and Jurkat lines, may be due to increased mTORC2 signaling required for oncogenic growth. However, a large number of these hits are also shared in common with HL60 where mTOR signaling is not particularly elevated. None of these hits have a strong impact on growth in K562 suggesting a distinct difference between bcr-abl mediated transformation and other modes of oncogenic transformation with regard to mTOR involvement.

### Different requirement for mTOR signaling in bcr-abl and non bcr-abl transformed cells

To investigate dependency on mTOR signaling, cells were titrated with rapamycin and growth measured after 3 days. OPM2 was highly sensitive to rapamycin while K562 was least affected (Fig 3A). HL60 and Jurkat cells had reduced proliferation that may be due to the general cytostatic effect of rapamycin on cell proliferation. However, reduced proliferation in OPM2 after rapamycin treatment appears greater than expected from growth arrest alone. Annexin V staining showed rapamycin induced apoptosis in OPM2 cells while having no effect on K562 (Fig 3B). Rapamycin only partially inhibits mTORC1 and is usually ineffective on mTORC2. We therefore used Torin1, an active site inhibitor of both mTORC1 and mTORC2 [8]. Torin1 had a strong negative effect on cell proliferation in HL60 and OPM2 and also decreased growth of Jurkat cells (Fig 3A). Inhibition of both TOR complexes however, had little effect on proliferation in K562.

**Fig 3.**
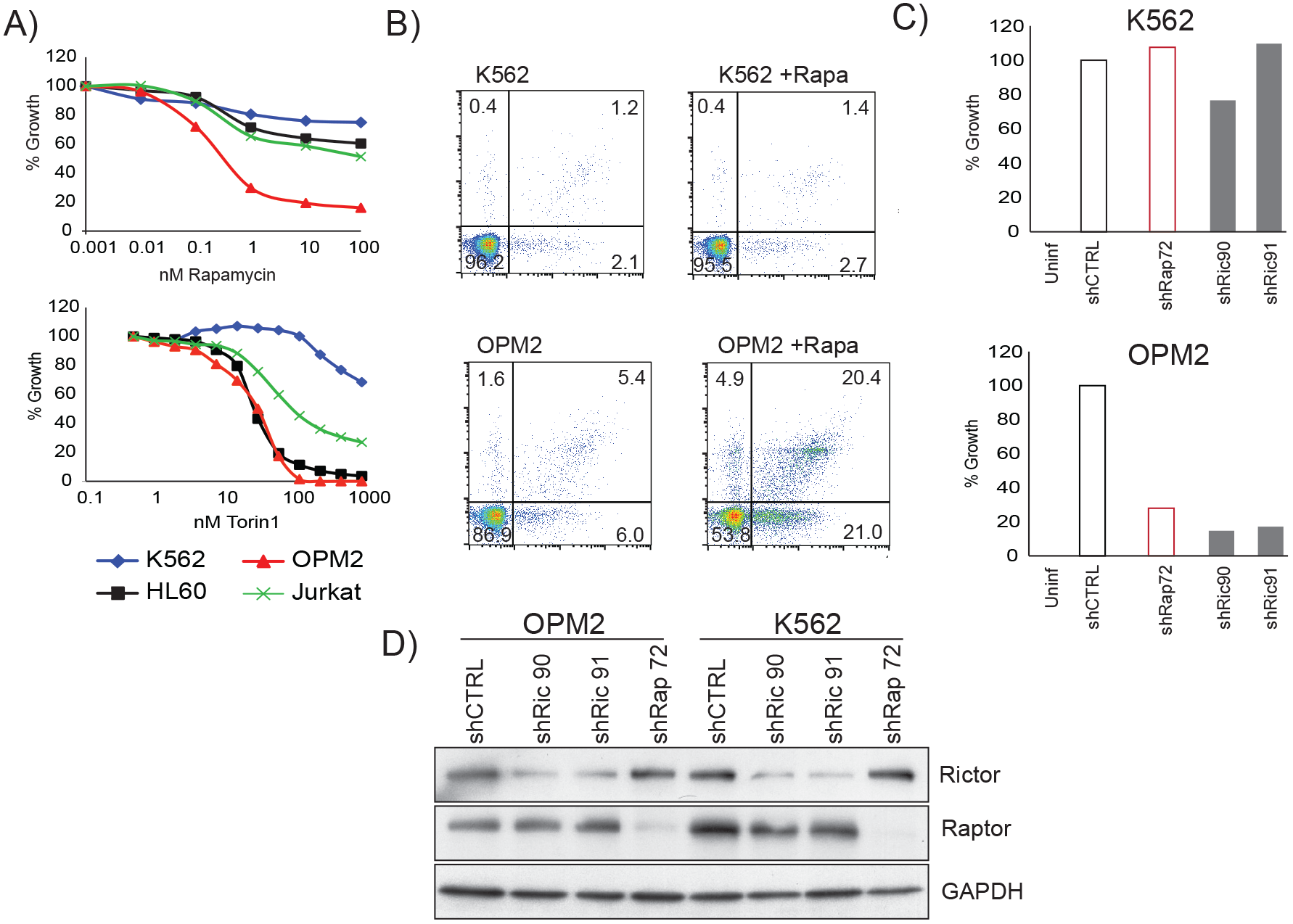
(A) Cell proliferation curves of the human cancer cell lines after treatment for 3 days with mTOR inhibitors. (B) FACS analysis of K562 and OPM2 cells stained for the apoptotic marker AnnexinV after treatment with rapamycin for 20 hours. (C) Proliferation of K562 and OPM2 cells 3 days after shRNA knockdown of Raptor and Rictor. (D) Immunoblot verifying Raptor and Rictor shRNA knockdown.

To further dissect the role of mTOR signaling, we performed shRNA knockdown of Rictor and Raptor on cell lines with the greatest divergence in response to mTOR inhibition, namely OPM2 and K562. shRNA knockdown of Raptor (mTORC1) or Rictor (mTORC2) was deleterious to OPM2 while having little effect on K562 (Fig 3C, 3D). Thus both mTORC1 and mTORC2 are essential for OPM2 growth and viability. In contrast mTOR signaling does not appear to be critical in K562, at least over the time-scale of the assay.

### PHD3 is essential for growth and survival in mTOR dependent OPM2

We wished to identify potential cancer therapeutic targets from the list of hits in Table S2. Of the essential genes in non bcr-abl transformed lines, only mTOR, Rictor, ATP1b3 and PHD3 can be pharmacologically inhibited. Rictor and mTOR signaling are already recognised as bona fide drug targets in oncology while inhibitors of ATP1b3 are non-specific and broadly target many Na^+^/K^+^ ATPase transporters. We focused on PHD3 (Prolyl hydroxylase 3, EGLN3) which is druggable and has been implicated in cancer [9].

A growth assay for cells treated with the general prolyl hydroxylase inhibitor DMOG (Fig 4A) showed that OPM2 was the most sensitive and K562 the least sensitive cell line. Thus response to DMOG within the cell panel mirrors sensitivity to the mTOR inhibitors rapamycin and Torin1 (Fig 3A). OPM2 acutely treated with DMOG showed reduced phosphorylation of S6K but not 4EBP (Fig 4B). DMOG appears to selectively inhibit phosphorylation of mTORC1 substrates in a similar fashion as rapamycin [10].

**Fig 4.**
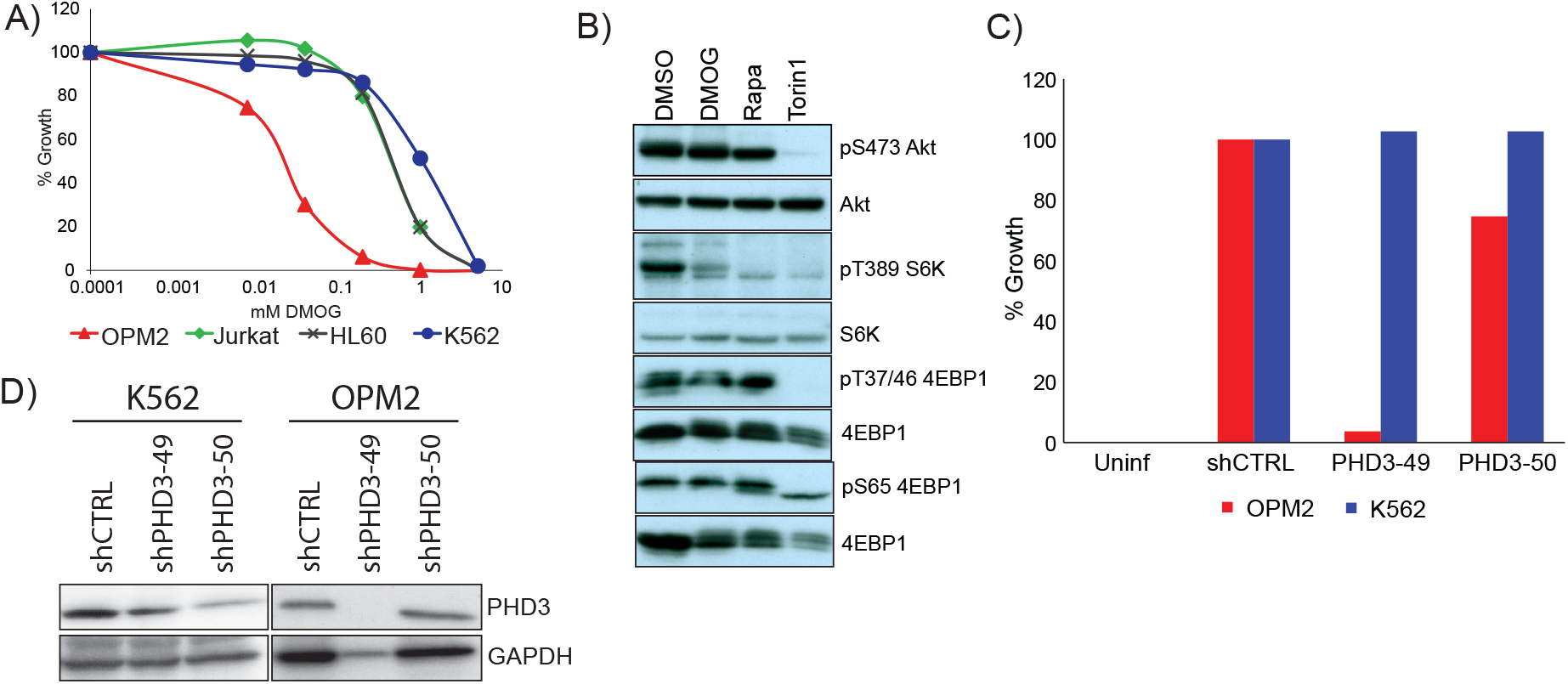
(A) Cell proliferation curves of the human cancer cell lines after treatment for 3 days with DMOG. (B) Immunoblot for mTOR activity read-outs in OPM2 after 1 hour drug treatment. (C) Proliferation of K562 and OPM2 cells after PHD3 shRNA knockdown. (D) Immunoblot verifying shRNA knockdown of PHD3.

DMOG inhibits all three cellular prolyl hydroxylases. To determine if the effect on growth was specific to PHD3 inhibition, OPM2 and K562 were infected with lentiviruses bearing shRNAs targeting PHD3. The degree of PHD3 knockdown correlated with growth suppression in OPM2 (Fig 4C, 4D). The great number of dying cells in OPM2 infected with shPHD3-49 reduced the amount of protein extracted for immunoblotting (Fig 4D, GAPDH loading control). In contrast, PHD3 knockdown had no effect on growth in K562.

## Discussion

Upregulation of mTOR signaling is a common event in cancer. mTORC1 promotes anabolic processes such as protein translation, lipid and nucleotide biosynthesis and suppresses catabolic processes like autophagy [11].

Under normal conditions this maintains the balance between anabolism and catabolism for homeostasis. However hyperactivation in cancer supports unrestricted growth due to elevated anabolic synthesis of cellular components. mTORC2 phosphorylates several members of the AGC kinase family. Among its downstream effects are regulation of cytoskeletal remodeling and migration, and via Akt (an important oncogene), it also promotes cell proliferation and survival. The full spectrum of mTOR’s role in the cell still remains to be characterised and there are presumably many more activities perturbed by mTOR dysregulation that are beneficial for cancer proliferation. Thus mTOR hyperactivation, even in cases where the driver mutation does not directly activate mTOR, is often recruited by oncogenic growth programs to support unrestricted proliferation.

The bcr-abl oncoprotein results from fusion of the abelson kinase gene, (ABL1) on chromosome 9, with the breakpoint cluster gene (BCR) on chromosome 22. The Abl kinase domain of bcr-abl is constitutively active due to loss of an autoinhibitory domain. Bcr-abl signaling causes chronic myelogenous leukemia and some cases of acute lymphoblastic leukemia or acute myelogenous leukemia [12]. Bcr-abl activates multiple downstream pathways (RAS/MAPK/ERK, JAK/STAT). The narrow range of histotypes generated by bcr-abl fusion however, indicates that the oncogenic program set in motion by bcr-abl signaling is restricted to cells from the hematopoietic lineage.

Our results show that genes essential for survival in an mTORC2 driven murine cancer model are not required in an isogenic bcr-abl driven cancer model (Fig 2A). This indicates that different signaling networks are activated downstream of either oncogenic driver (Pten loss vs. bcr-abl expression), and that these networks do not appear to share common effector proteins to execute the various essential functions.

This observation with the mouse model could be extended to human cancer cell lines. A substantial fraction of human counterparts of the genes previously validated in the murine model were also essential in a human cell line panel with the exception of bcr-abl transformed K562. In contrast to the other cell lines (OPM2, Jurkat, HL60), K562 was insensitive to pharmacological mTOR inhibition and shRNA knockdown of mTOR complexes. We could thus confirm a clear divide between the suite of essential genes in an mTOR dependent vs. bcr-abl transformed phenotype. The cell lines were initially selected without consideration for mTOR dependency. However the central importance of mTOR in coordinating cell growth and metabolism with nutrient and growth factor availability, results in the recruitment of mTOR hyperactivation regardless of the original oncogenic lesion. Thus there may be a general dependency for mTOR signaling to be recruited in diverse oncogenic programs (with bcr-abl oncogenic signaling being an exception).

The validated hits are potential cancer therapeutic targets. Among druggable targets, we focused on PHD3. While generally regarded as a tumor suppressor, PHD3 overexpression is a marker for poor prognosis in clear-cell renal cell carcinoma (ccRCC) [13], one of the few cancers to respond to rapamycin treatment [14] and supports translation and glycolysis in ccRCC [15]. Depletion of PHD3 activity by pharmacological inhibition or shRNA knockdown led to growth reduction and decreased mTORC1 signaling in OPM2 (Fig 4B, 4C). mTORC1 activation was previously shown to be sensitive to PHD inhibition [16]. Thus PHD3 is essential for cell viability in a transformed cell line where mTOR signaling is involved in driving cell growth and proliferation.

## Acknowledgements

This work was funded by the Krebsliga beider Basel.

## Materials and Methods

### Cell culture

All cells were grown in Iscove’s medium supplemented with 10% FCS, 2mM L-glutamine, 100U/ml penicillin, 100μg/ml streptomycin and 50μM b-mercaptoethanol at 37°, 5% CO_2_. Derivation of 15V4, 6.5, 6.8 and shP cells has been previously described [3]. b-abl cells were obtained by expressing a bcr-abl construct in 15V4 and selecting for IL3 independence [6]. Murine mast cell medium was supplemented with mouse IL3 from conditioned medium as previously described [3].

### Cell proliferation assays

Cells were plated in 96-well microtitre plates at a density of 3000 cells/150μl medium and growth measured after 3 days by addition of 0.1vol resazurin (final concentration 50μM). Fluorescence was read at 535/595nm (Ex/Em) with a plate reader.

### Immunoblotting

Cells were lysed in RIPA buffer (containing protease and phosphatase inhibitors) and total protein (20μg) loaded on SDS-PAGE gels. Proteins were blotted onto nitrocellulose membranes and probed with commercially available antibodies against pS6K T389 (#9205), pAkt S473 (#9271), PTEN (#9559), total S6K (#9202), total Akt (#9272), Raptor (#2280), Rictor (#2114) from Cell Signaling, PHD3 (Thermo Fisher PA1-16526) and GAPDH (Abcam ab9385). Signals were detected with HRP-conjugated secondary antibodies and visualized with ECL detection reagent (GE Healthcare).

### shRNA knockdown screens

MISSION TRC shRNA lentiviral library containing 80’000 lentiviral clones targeting 15’000 genes was purchased from Sigma-Aldrich. Cells (3,000 per well) were infected with individual lentiviruses (at a m.o.i. of ~1) in separate wells using a robotic platform. After 24h infection, cells were plated either with/without IL3 and puromycin (0.8 μg/ml) added to kill non-infected cells. On day 5, relative growth (compared to infection with shControl virus) was quantified by measuring cell proliferation with resazurin, and Growth_+IL3_/Growth_−IL3_ was used to calculate a ratio of inhibition (RI). The RI was calculated using the mean of the two best performing shRNA constructs per gene (there are usually 5 shRNAs per gene).

Similar steps were carried out for the human shRNA screen, except for omission of a parallel +IL3 screen. Actual lentiviral infectivity was determined by measuring acquired puromycin resistance in murine bcr-abl cells (assuming that shRNAs targeting the human gene would be ineffective on their murine homologs). This allowed normalization of viral titers across all the lentiviruses used. The ratio of inhibition for a particular lentivus was calculated by dividing (proliferation in infected human cells)/(proliferation in murine b-abl cells). Lentiviruses targeting the essential ribosomal genes RPL32 and RPS17 were used as positive controls for cell death. The ratio of inhibition for RPS17 was set at 100% and all other ratios expressed as a percentage of this.

### shRNA lentiviruses

Lentiviral article were ordered from Sigma-Aldrich. Human Raptor (TRCN0000039772), human Rictor (TRCN0000074290, TRCN0000289691), human PHD3 (TRCN0000001049, TRCN0000001050). Cells were infected overnight

### FACS analysis

Cells were stained with Annexin V (Invitrogen) and counter stained with propidium iodide (5μg/ml) as a vital stain. Apoptotic cells detected by FACS analysis in the FL-1 channel and living cells were simultaneously detected in a separate channel.

## Supplementary Data

**Table S1.** Ratio of inhibition for the 25 top-scoring genes in 3 mTORC2-addicted lines (6.5, 6.8, shP), and in b-abl cells. RI values for each gene using the mean of two best scoring oligos for growth^−IL3^/growth^+IL3^ were calculated as described [3]. An RI value >5 corresponds to inhibition of proliferation by >80%.

| Gene symbol | NM id | Gene name | Mean of Two 6.5 | Mean of Two 6.8 | Mean of Two shP | Mean of Two b-abl |
| --- | --- | --- | --- | --- | --- | --- |
| **Abca3** | NM_013855 | ATP-binding cassette | 22.5 | 33.3 | 30.4 | 3.6 |
| **Aqp1** | NM_007472 | aquaporin 1 | 10.3 | 30.7 | 6.4 | 0.8 |
| **Aqp7** | NM_007473 | aquaporin 7 | 12.7 | 64.7 | 23.1 | 1.2 |
| **Arsb** | NM_009712 | arylsulfatase B | 16.3 | 19.4 | 26.8 | 0.9 |
| **Asb15** | NM_080847 | ankyrin repeat and SOCS box-containing protein 15 | 11.0 | 65.6 | 8.5 | 0.7 |
| **Atp1b3** | NM_007502 | ATPase | 20.1 | 33.3 | 15.2 | 3.7 |
| **Cbx2** | NM_007623 | chromobox homolog 2 (Drosophila Pc class) | 10.2 | 8.0 | 7.2 | 1.9 |
| **Cox7a2l** | NM_009187 | cytochrome c oxidase subunit VIIa polypeptide 2-like | 23.2 | 12.9 | 26.9 | 2.8 |
| **Dmgdh** | NM_028772 | dimethylglycine dehydrogenase precursor | 13.9 | 23.5 | 17.3 | 1.8 |
| **Fhl3** | NM_010213 | four and a half LIM domains 3 | 23.2 | 63.6 | 34.6 | 3.6 |
| **Il1f5** | NM_019451 | interleukin 1 family | 23.8 | 86.7 | 16.1 | 1.5 |
| **Kctd11** | NM_153143 | potassium channel tetramerisation domain containing 11 | 7.4 | 29.4 | 5.4 | 1.3 |
| **Mff** | NM_029409 | Mitochondrial fission factor | 10.6 | 32.1 | 6.4 | 1.3 |
| **Mmd2** | NM_175217 | monocyte to macrophage differentiation-associated 2 | 9.0 | 16.6 | 13.2 | 1.8 |
| **Mrpl27** | NM_053161 | mitochondrial ribosomal protein L27 | 19.2 | 24.8 | 28.3 | 1.0 |
| **mTOR** | NM_020009 | mammalian Target Of Rapamycin | 24.5 | 69.7 | 64.9 | 0.8 |
| **Phykpl** | NM_028398 | 5-phosphohydroxy-L-lysine phospho-lyase | 12.9 | 13.3 | 9.6 | 1.3 |
| **Prdm4** | NM_181650 | PR domain containing 4 | 34.2 | 89.0 | 55.7 | 3.9 |
| **Phd3** | NM_028133 | Prolyl hydroxylase 3 | 12.8 | 14.9 | 9.2 | 1.1 |
| **Rab32** | NM_026405 | RAB32 | 16.3 | 48.0 | 13.1 | 1.1 |
| **Rab33a** | NM_011228 | RAB33A | 7.9 | 46.2 | 26.8 | 1.2 |
| **Rictor** | NM_030168 | Rictor | 29.7 | 76.4 | 10.3 | 0.7 |
| **S100a11** | NM_016740 | S100 calcium binding protein A11 | 9.8 | 11.3 | 24.9 | 1.4 |
| **Sfn** | NM_018754 | stratifin | 22.7 | 30.4 | 23.6 | 1.3 |
| **Sh3bp2** | NM_011893 | SH3-domain binding protein 2 | 8.8 | 40.8 | 5.8 | 1.2 |

**Table S2.**
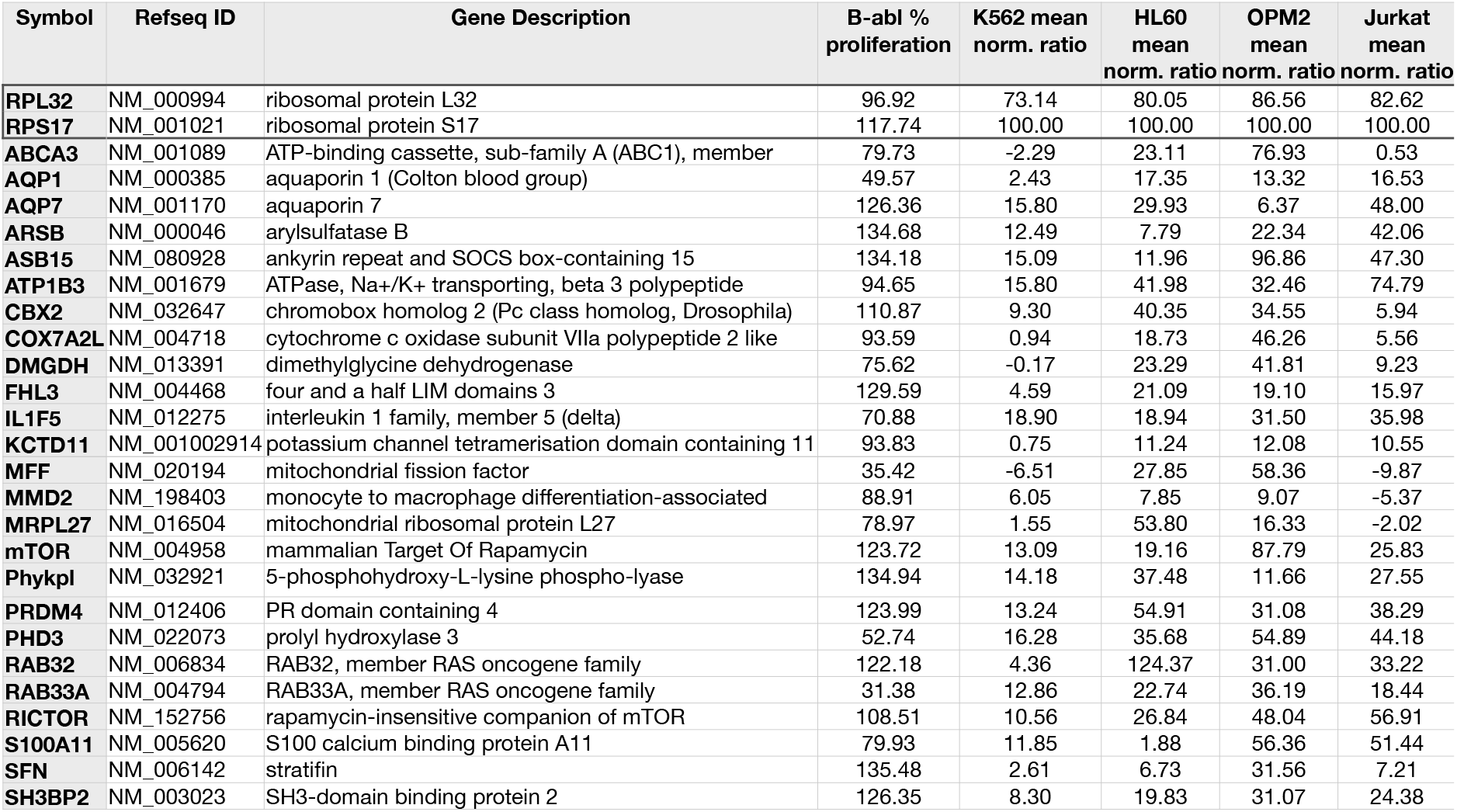
Ratio of inhibition for the human homologs of the 25 genes from Table S1 tested on a panel of human cancer cell lines. Essential ribosomal genes RPL32 and RPS17 were included as positive controls for cell death. RI for each shRNA was normalized to RPS17.

## References

[1] J. Kim, K.-L. Guan, mTOR as a central hub of nutrient signalling and cell growth, Nat. Cell Biol. 21 (2019) 63–71. doi:10.1038/s41556-018-0205-1.

[2] E. Rad, J.T. Murray, A.R. Tee, Oncogenic Signalling through Mechanistic Target of Rapamycin (mTOR): A Driver of Metabolic Transformation and Cancer Progression, Cancers (Basel). 10 (2018) 5. doi:10.3390/cancers10010005.

[3] M. Colombi, K.D. Molle, D. Benjamin, K. Rattenbacher-Kiser, C. Schaefer, C. Betz, et al., Genome-wide shRNA screen reveals increased mitochondrial dependence upon mTORC2 addiction, Oncogene. 30 (2011) 1551–1565. doi:10.1038/onc.2010.539.

[4] K.F. Kiser, M. Colombi, C. Moroni, Isolation and characterization of dominant and recessive IL-3-independent hematopoietic transformants, Oncogene. 25 (2006) 6595–6603. doi:10.1038/sj.onc.1209673.

[5] E. Andrejauskas, C. Moroni, Reversible abrogation of IL-3 dependence by an inducible H-ras oncogene, Embo J. 8 (1989) 2575–2581.

[6] L. Wyder Peters, K.D. Molle, A. Thiemeyer, A. Knopf, M. Goxe, P. Guerry, et al., An isogenic cell panel identifies compounds that inhibit proliferation of mTOR-pathway addicted cells by different mechanisms, J Biomol Screen. 19 (2014) 131–144. doi:10.1177/1087057113497798.

[7] T.R. Breitman, S.J. Collins, B.R. Keene, Replacement of serum by insulin and transferrin supports growth and differentiation of the human promyelocytic cell line, HL-60, Exp. Cell Res. 126 (1980) 494–498.

[8] C.C. Thoreen, S.A. Kang, J.W. Chang, Q. Liu, J. Zhang, Y. Gao, et al., An ATP-competitive mammalian target of rapamycin inhibitor reveals rapamycin-resistant functions of mTORC1, J. Biol. Chem. 284 (2009) 8023–8032. doi:10.1074/jbc.M900301200.

[9] Q.-L. Liu, Q.-L. Liang, Z.-Y. Li, Y. Zhou, W.-T. Ou, Z.-G. Huang, Function and expression of prolyl hydroxylase 3 in cancers, Arch Med Sci. 9 (2013) 589–593. doi:10.5114/aoms.2013.36987.

[10] A.Y. Choo, S.-O. Yoon, S.G. Kim, P.P. Roux, J. Blenis, Rapamycin differentially inhibits S6Ks and 4E-BP1 to mediate cell-type-specific repression of mRNA translation, Proc. Natl. Acad. Sci. U.S.a. 105 (2008) 17414–17419. doi:10.1073/pnas.0809136105.

[11] R.A. Saxton, D.M. Sabatini, mTOR Signaling in Growth, Metabolism, and Disease, Cell. 169 (2017) 361–371. doi:10.1016/j.cell.2017.03.035.

[12] R. Kurzrock, H.M. Kantarjian, B.J. Druker, M. Talpaz, Philadelphia chromosome-positive leukemias: from basic mechanisms to molecular therapeutics, Ann. Intern. Med. 138 (2003) 819–830.

[13] J. Garcia-Donas, L.J. Leandro-García, A. González Del Alba, M. Morente, I. Alemany, E. Esteban, et al., Prospective study assessing hypoxia-related proteins as markers for the outcome of treatment with sunitinib in advanced clear-cell renal cell carcinoma, Ann. Oncol. 24 (2013) 2409–2414. doi:10.1093/annonc/mdt219.

[14] C. Battelli, D.C. Cho, mTOR inhibitors in renal cell carcinoma, Therapy. 8 (2011) 359–367. doi:10.2217/thy.11.32.

[15] P. Miikkulainen, H. Högel, K. Rantanen, T. Suomi, P. Kouvonen, L.L. Elo, et al., HIF prolyl hydroxylase PHD3 regulates translational machinery and glucose metabolism in clear cell renal cell carcinoma, Cancer Metab. 5 (2017) 5. doi:10.1186/s40170-017-0167-y.

[16] R.V. Durán, E.D. MacKenzie, H. Boulahbel, C. Frezza, L. Heiserich, S. Tardito, et al., HIF-independent role of prolyl hydroxylases in the cellular response to amino acids, Oncogene. 32 (2013) 4549–4556. doi:10.1038/onc.2012.465.

